# Cortical circuit-based lossless neural integrator for perceptual decision-making

**DOI:** 10.1101/158865

**Authors:** Jung H. Lee, Joji Tsunada, Sujith Vijayan, Yale E. Cohen

## Abstract

The intrinsic uncertainty of sensory information (i.e., evidence) does not necessarily deter an observer from making a reliable decision. Indeed, uncertainty can be reduced by integrating (accumulating) incoming sensory evidence. It is widely thought that this accumulation is instantiated via recurrent rate-code neural networks. Yet, these networks do not fully explain important aspects of perceptual decision-making, such as a subject’s ability to retain accumulated evidence during temporal gaps in the sensory evidence. Here, we utilized computational models to show that cortical circuits can switch flexibly between ‘retention’ and ‘integration’ modes during perceptual decision-making. Further, we found that, depending on how the sensory evidence was readout, we could simulate ‘stepping’ and ‘ramping’ activity patterns, which may be analogous to those seen in different studies of decision-making in the primate parietal cortex. This finding may reconcile these previous empirical studies because it suggests these two activity patterns emerge from the same mechanism.

## Introduction

One of the fundamental operations of the brain is to transform representations of external sensory stimuli (i.e., sensory evidence) into a categorical judgment, despite the inherent uncertainty of this sensory evidence. For instance, we can determine the direction of the wind, even though its instantaneous direction continuously fluctuates. It is widely thought that this moment-by-moment uncertainty is minimized by temporally integrating (accumulating) this incoming sensory evidence^1–4^. Potential neural correlates of this accumulation process have been identified in a variety of brain areas, including the lateral intraparietal cortex (area LIP)^2, 3, 5^, the prefrontal cortex^6^, and the frontal eye fields^7^. In particular, spiking activity in these brain areas appears to smoothly ‘ramp up’ (accumulate; i.e. linearly increasing activity over time) prior to a perceptual decision. Further, the rate of this accumulation, which governs the time to reach a decision threshold (i.e., the time to the perceptual decision), is correlated with the ambiguity of the sensory evidence: as the evidence becomes less ambiguous (e.g., the instantaneous fluctuations in wind direction decrease), the rate of the ramping increases^3^.

Such neural integration has been modeled in two very different ways, each of which relies on different coding strategies and mechanisms of integration^1^. In the first type of model, rate-code neural integrators (NI) integrate sensory evidence and represent accumulated evidence as monotonically increasing (‘ramping’) spiking activity. In this rate-code model, the firing rates of individual neurons increase over time in response to continuous inputs^2, 3, 8^. In an alternative model, location-code NIs store accumulated evidence as the location of highly elevated spiking activity. In such a location-code NI, the location of these highly active neurons, which are referred to as a ‘bump’, travels through a network over time^9, 10^. That is, the location of bump activity corresponds to the total amount of accumulated evidence.

Because ramping activity has been found in several studies of perceptual decision-making^1, 3^, it is generally believed that a rate-code NI is the natural circuit candidate for neural integration of sensory information. However, recent behavioral studies have questioned whether a rate-code NI can, in fact, accurately capture the dynamics of perceptual decision-making. For example, a temporal gap between stimulus presentations has little impact on the accuracy of an observer’s behavioral choices^11, 12^, indicating that accumulated evidence can be maintained during this temporal gap. Yet, during this gap, the firing rates of neurons in a rate-code NI are likely to deviate from the desired values if the network is perturbed even slightly^11^. This deviation can occur because a rate-code NI’s feedback (recurrent) inputs and its leaky currents have to be precisely balanced in order to maintain the desired values during such temporal gaps^11, 13^.

Further, the nature of neuronal activity during decision-making calls into question the suitability of rate-code NIs. Traditionally, as noted above, decision-making activity, at both the single-neuronal and population level, was thought to be best described as ramping activity^2, 3, 5^. However, recent studies indicate that, whereas population-level activity can be thought of as ramping, single-neuronal activity may be better described as discrete ‘steps’ (changes) in neuronal activity^14–16^. If this stepping activity is an accurate descriptor of decision-making activity, it follows that rate-code NIs may not be an appropriate model: because single-neuron activity and population-level activity in rate-code NI are completely correlated in rate-code NIs, it is not clear how single-neuron stepping activity can become population-level ramping activity.

In contrast to a rate-code NI, a location-code NI can maintain stable states even in the absence of external inputs^9^. Further, neurophysiological studies have identified sequential activation that is similar to this propagation of bump activity^17–21^, and a theoretical study^22, 23^ proposed that such a network can be constructed out of commonly found depressing synapses^24–29^. Inspired by these findings, we constructed a computational model of cortical circuits with depressing synapses to test the hypothesis that a location-code NI is a viable model for perceptual decision-making.

We found that, like previous location-code NIs, a neurobiologically inspired network can sustain bump activity at a specific location when there is a temporal gap in sensory evidence, whereas sensory evidence causes bump activity to propagate through the network. Our model is unique in that it is based on depressing synapses and the interplay between two commonly found inhibitory neuron types^30, 31^. We also found that the sensory evidence, which is stored as the location of bump activity, could be readout in two different modes, depending on the connections with downstream readout neurons. When the connectivity between integrator and downstream readout neurons was dense, readout neurons predominantly showed classic ramping activity as the sensory evidence was accumulated into a decision variable^1, 3, 5^. In contrast, when the connectivity was sparse, readout neurons predominantly exhibited stepping activity^16^; that is, the firing rate of individual neurons changed from one state to another transiently, whereas population activity gradually ramped over time. This observation predicts that either ramping or stepping modes can emerge, depending on the connectivity. This dual-readout mode may, in part, reconcile the degree to which components of decision-making are encoded as ramping- or stepping-like spiking activity.

## Results

This section describes how cortical circuits can implement a lossless integrator. The first subsection describes simulation results suggesting that generic cortical circuits (Fig. 1A), which contain two common types of inhibitory neurons and depressing synapses, can readily realize a lossless (‘perfect’) location-code NI. The second subsection discusses bifurcation analyses of abstract models of rate- and location-code NIs, which were conducted to examine how reliably these two types of NIs can retain sensory evidence during temporal gaps in the sensory evidence. In the third subsection, we propose a location-code NI that can have continuous attractors (Fig. 1B). Finally, we discuss how evidence accumulated in our integrators can be readout by downstream neurons. Interestingly, this readout activity maps onto two different modes of spiking activity that have been identified during neurophysiological studies of decision-making: classic ‘ramping’ activity^2^ and newly identified ‘stepping’ activity^16^.

**Figure 1:**
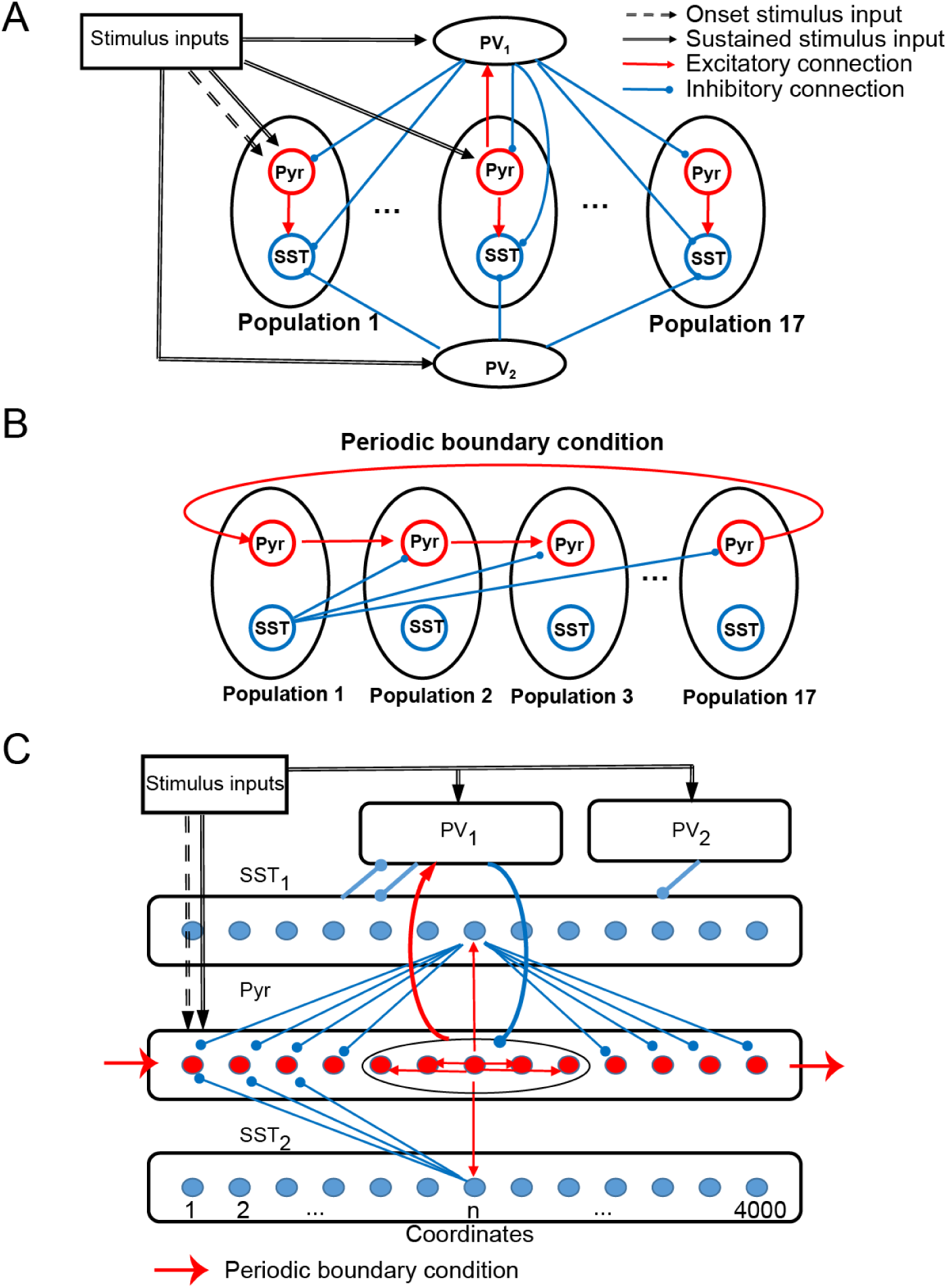
The structure of the two versions of our integrator. **(A)**, Connectivity between all 19 neuronal populations in the *discrete* integrator. **(B)**, Interconnectivity between the 17 Pyr-SST populations; see **Methods** and Tables 1 and 2 for more details and parameters. Red and blue arrows indicate excitatory and inhibitory connections within the network model, respectively. Dashed and thick black arrows represent onset and sustained stimulus inputs, respectively. **(C)**, Structure of *continuous* integrator. The five neuronal populations (Pyr, PV_1_, PV_2_, SST_1_, and SST_2_) interact with each other via connections shown in the figure. The thin red arrows and blue arrows represent the excitatory and inhibitory connections between individual neurons, respectively. In contrast, the thick arrows (including red and blue) show connections between the neuronal populations. All connections between populations are randomly established. Sensory inputs are introduced to Pyr, PV_1_ and PV_2_ (dashed arrows). Periodic boundary condition is used to connect Pyr cells, as shown in the red arrow; see **Methods** and Table 3 for more details and parameters.

**Table 1:** Neural parameters for neurons and synapses. When a spike arrived, the membrane potential instantly jumped to a new value, which was determined by its capacitance (C) and time constant (τ_m_). When the membrane potential was higher than the spike threshold, the membrane potential was reset (V_reset_). Without any external input, the membrane potential relaxed back its the resting membrane potentials (E_L_). Synaptic events decayed exponentially with a 2-ms time constant (τ_syn_). All synapses had a 1.5-ms delay unless otherwise stated; the only exception is given in Table 2. For depressing synapses, we selected the parameters (U and τ_ref_) given below.

| Neuronal Parameters |  | Synaptic parameters |  |
| --- | --- | --- | --- |
| C<br>(membrane capacitance) | 1 pF | $\tau_{\text{syn}}$ | 2.0 ms |
| $V_{\text{th}}$<br>(spike threshold) | 20 mV | delay | 1.5 |
| $\tau_m$<br>(Membrane time constant) | 20 ms | U | 0.2 |
| $E_L$<br>(resting membrane potential) | 0 mV | $\tau_{\text{ref}}$ | 200 ms for discrete integrator<br>500 ms for continuous integrator |
| $V_{\text{reset}}$<br>(reset after spiking) | 0 mV | | |

**Table 2:** The parameters of the discrete integrator. We connected populations by specifying connection probabilities and synaptic connection strengths. The first value in the parentheses is the connection probability. The connection strengths followed Gaussian distributions. The mean values of these distributions are the second value in the parentheses, and the standard deviations were 10% of the mean. The excitatory and inhibitory connections could not be less than or greater than 0, respectively; when they violated this condition, we set them to 0.

|  | Total Number | Background inputs<br>(Hz) | Stimulus input (Hz;<br>sustained) |
| --- | --- | --- | --- |
| Pyr | 6800 | 2,800 | 2000 |
| PV<sub>1</sub> | 1088 | 4,500 | 2000 |
| PV<sub>2</sub> | 1088 | N/A | 2000 |
| SST | 544 | 3,200 | N/A |
| Connectivity within populations (connection probability, strength in pA) |  |  |  |
| Pyr→Pyr | (1.0, 1.8) | Pyr→SST | (0.4, 0.96) |
| PV<sub>1</sub>→PV<sub>1</sub> | (0.3, -0.72) | PV<sub>1</sub>→PV<sub>1</sub> | (0.1, -0.72) |
| Connectivity across populations (connection probability, strength in pA) |  |  |  |
| Pyr→Pyr | (0.2, 0.12) *delay 10 ms | PV<sub>2</sub>→SST | (1.0, -6.0) |
| Pyr→PV<sub>1</sub> | (0.2, 0.12) | SST→Pyr | (1.0, -4.8) |
| PV<sub>1</sub>→Pyr | (0.2, -1.08) | SST→PV<sub>1</sub> | (0.3, -0.6) |
| PV<sub>1</sub>→SST | (0.3, -0.6) |  |  |
| Connection strength for background and stimulus inputs in pA |  |  |  |
| Pyr | 0.12 | PV<sub>2</sub> | 0.36 |
| PV<sub>1</sub> | 0.12 | SST | 0.12 |
| Onset stimulus input |  |  |  |
| Target | Pyr neurons<br>in population 1 | Firing rate | 1000 Hz |

**Table 3:** The parameters of the continuous integrator. Due to the lack of population structure, we connected neurons by specifying the number of presynaptic neurons to each neuron type. The frequency of stimulus inputs given below is the default value used unless stated otherwise; see also Equation 3. The first value is the number of presynaptic neurons, and the second value is the connection strength in pA. The excitatory and inhibitory connections could not be less than or greater than 0, respectively; when they violated this condition, we set them to 0. The background inputs to all neurons in the continuous integrator are mediated by synapses whose strength are 0.13 pA.

|  | Total Number | Background inputs (Hz) | Stimulus input (Hz) |
| --- | --- | --- | --- |
| Pyr | 4000 | 3,850 | 4,800 |
| PV<sub>1</sub> | 1000 | 3,850 | 1,200 |
| PV<sub>2</sub> | 1000 | 3,000 | 1,200 |
| SST<sub>1</sub> | 4000 | 2,000 | N/A |
| SST<sub>2</sub> | 4000 | 2,000 | N/A |
| Connectivity (Number of presynaptic neurons, strength in pA) |  |  |  |
| Pyr→Pyr | (400, 0.52) | PV<sub>1</sub>→SST<sub>1</sub> | (150, -0.78) |
| Pyr→PV<sub>1</sub> | (400, 0.52) | PV<sub>2</sub>→SST<sub>1</sub> | (1000, -0.78) |
| Pyr→SST<sub>1</sub> | (1, 11.7) | SST<sub>1</sub>→Pyr | (3600, -0.78) |
| Pyr→SST<sub>2</sub> | (1, 11.7) | SST<sub>1</sub>→PV<sub>1</sub> | (1200, -0.78) |
| PV<sub>1</sub>→Pyr | (160, -1.87) | SST<sub>2</sub>→Pyr | (400, -0.78) |
| PV<sub>1</sub>→PV<sub>1</sub> | (160, -0.78) |  |  |

### Cortical circuits can readily implement lossless location integrator

Cortical circuits have three common properties that are relevant for our model. First, pyramidal (Pyr) neurons in sensory cortex are topographically organized as a function of their sensory response profiles via spatial^32, 33^ and functional^34^ connections. Second, cortical circuits also contain parvalbumin positive (PV) and somatostatin positive (SST) inhibitory interneurons^30^. PV neurons have a fast-spiking pattern of activity, whereas SST neurons have a low-threshold spiking pattern. For our purposes, it is important to note that, although most inhibitory interneurons are broadly tuned to sensory inputs, the response profiles of SST neurons can be as sharply tuned as those of Pyr neurons^35^. Third, via lateral inhibition, SST neurons inhibit neighboring cortical neurons^36–39^.

Based on previous modelling studies^22, 23^ that proposed propagating bump activity can be elicited by depressing synapses, we built a cortical network model (Fig. 1A), in which Pyr neurons interacted with one another through intra-population depressing synapses^24–29^ and inter-population unidirectional static synapses. We refer to this cortical network model as the ‘discrete’ integrator; see Methods for more details. Transient sensory stimuli (100 ms), which mimicked sensory-driven onset responses in sensory cortex^40–43^, only drove Pyr cells in the first population. In contrast, sustained sensory stimuli (after 100 ms) drove Pyr neurons in all neuronal populations. In our first simulation, we only provided Pyr and PV neurons with sensory evidence at two discrete time intervals: time=100-300 ms and during time=800-1000 ms.

As seen in Fig. 2A, the Pyr populations were sequentially activated by sensory stimulation. Further, on average, both populations of PV neurons were more active during sensory stimulation than during the temporal gap (Fig. 2B). More importantly, when there was a temporal gap in the sensory evidence (as indicated by the black double-headed arrow in Fig. 2A), the sequential activation of the network stopped but activity was maintained by a specific population of Pyr neurons (Pyr population 5 in Fig. 2A). That is, during a temporal gap in the sensory evidence, the network retained the accumulated information, a finding that is consistent with lossless integration. When we presented the second sensory stimulus, information resumed propagating through the network as seen by the sequential activation of Pyr population 6, followed by population 7, etc.

**Figure 2:**
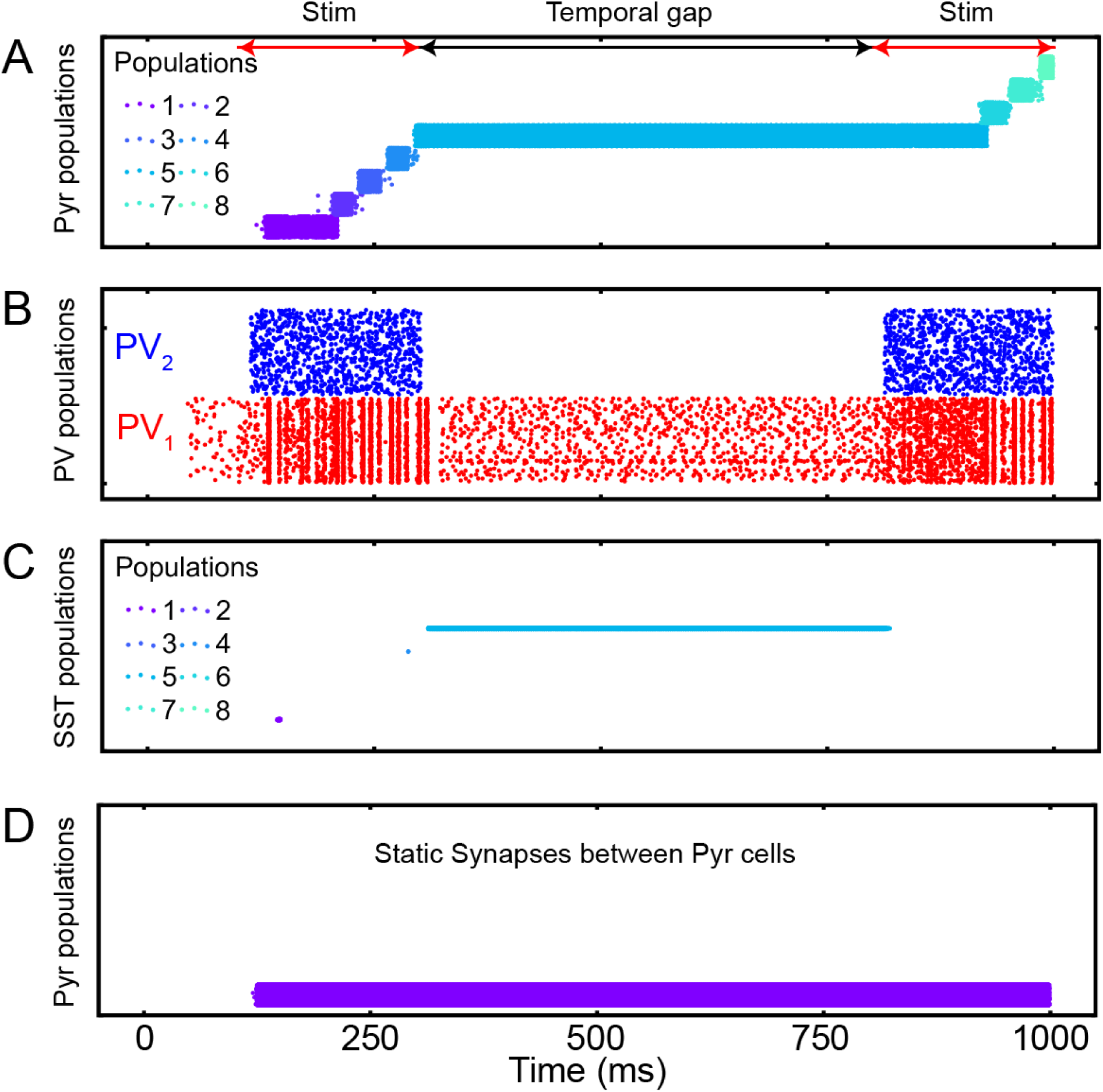
The responses of populations of the discrete integrator. **(A)**, Spiking activity of Pyr neurons in all 17 neuronal populations; each population had 400 Pyr neurons. Each row in the plot shows the spike times of an individual Pyr neuron. Each of the 17 populations are shown in a different color; see legend for the color codes of a subset of these populations. The red and black arrows show sensory-stimulus periods and the temporal gap between them, respectively. During a 1000 msec-long simulation, we noted that only 8 populations were activated. **(B)**, PV_1_ and PV_2_ activity during the sensory-stimulus periods and the temporal gap between both. Both PV populations contained 1088 PV neurons. **(C)**, SST neuron activity in all 17 populations; there are 16 SST neurons in each population. The same color scheme is used as in (**A**). SST neurons became active only during the temporal gap, and they belong to the same population. **(D)** Pyr activity when all depressing synapses are replaced with static ones.

When we explored the network in more detail, we found key roles for the inhibitory neurons and for the depressing synapses. For example, SST neurons were active only during the temporal gap (Fig. 2C) and that bump activity did not propagate when we replaced the depressing synapses with static synapses (Fig. 2D). We also noted that the non-specific feedback inhibition of PV_1_ neurons play a key role to activate an appropriate population of neurons (i.e., Pyr population 6 in Fig. 2A, following the temporal gap). Without this inhibition, when we presented the second sensory stimulus, Pyr population 1 (which was activated by the first initial 100-ms of sensory stimulation) was inappropriately activated. This altered the amount of accumulated information (supplemental Fig. 1).

### The stability of sensory evidence during a temporal gap in location-code NIs and in rate-code NIs

Next, we asked whether a location-code NI could retain sensory evidence during a temporal gap more reliably than a rate-code NI. To address this question, we created close-form firing-rate models that described the rate- and location-code NIs. We modeled a rate-code NI with a single recurrent neural population^1^ (Equation 1; see the inset of Fig. 3A), whereas we modeled a location- code NI with two recurrent neural populations because it relies on the sequential activation of neurons (Equation 2).

**Figure 3:**
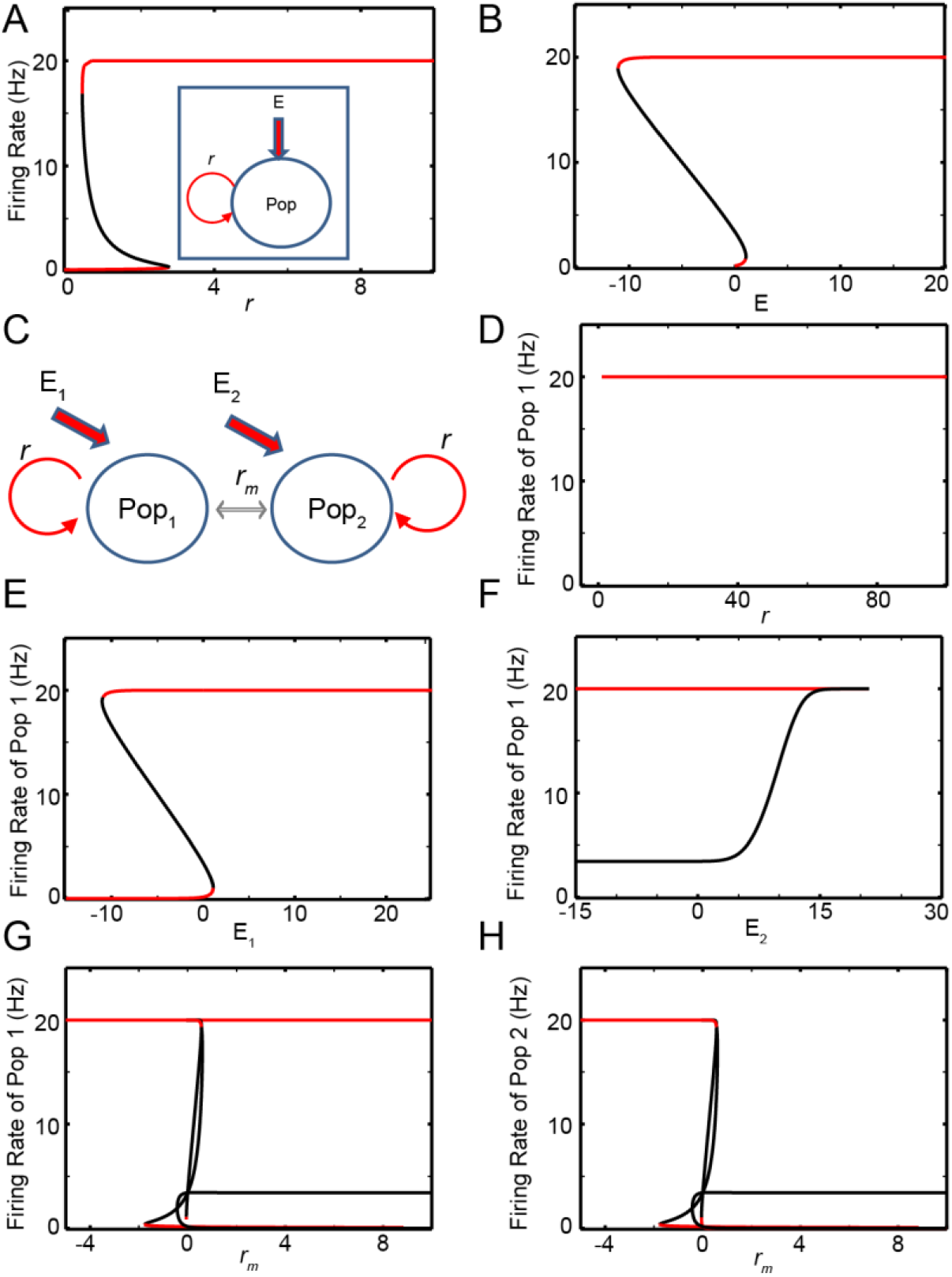
The bifurcation analysis of rate- and location-code NIs. **(A)** and **(B)**, Bifurcation analyses with the recurrent connections (*r*) and the external inputs (*E*) as bifurcation parameters for the recurrent rate-code network model, respectively; the schematics this network model is shown in the inset of (**A**). (**C**), Schematic of the reduced model of location-code NI. **(D)**-**(F)** Bifurcation analysis of the firing rate of population 1 with respect to within-population recurrent connections (*r*) and external input to populations 1 and 2 (*E*_1_, *E*_2_), respectively. **(G)** and **(H)**, Bifurcation analysis of firing rate of populations 1 and 2, respectively, in terms of the lateral interactions (*r*_m_). Red and black lines represent stable and unstable steady solutions, respectively. Pop: neuronal population.

The firing rate of the rate-code recurrent network obeys Equation 1^1^:

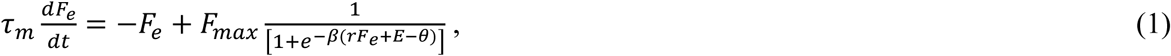

where *F*_e_ and *r* are the firing rate and recurrent connection strength, respectively; *F*_max_ is the maximum firing rate; *θ* is the spiking threshold; *E* is the external input; and *β* represents the strength of stochastic inputs^44^. *F*_e_ represents the leak current. The selected default parameters are *F*_max_=20, *β=*1*, θ*=0.5, *r*=1 and *E*=0, unless stated otherwise. We modeled the gain (transfer function; i.e., the number of spikes that a neuron can generate in response to afferent synaptic activity) with a logistic function^44^. The firing rate of this neuron increases, as *r* increases, which represents the relative strength of recurrent inputs; in our model, *r* is dimensionless.

We tested the stability of this rate-code NI during a temporal gap, in which external inputs are absent (i.e., *E*=0), by conducting a bifurcation analysis with the XPPAUT analysis platform^45^. A bifurcation analysis identifies the steady-state solutions, in which a system can stay indefinitely until perturbed. Moreover, this analysis clarifies whether the steady-state solutions are stable in response to perturbations of the bifurcation parameters (which, in our analysis, is either the strength of the recurrent connections *r* or the external inputs *E*; see the inset of Fig. 3A). That is, we tested if a rate-code NI is stable in response to small changes in either recurrent inputs or external inputs.

In Figs. 3A and B, the stable and unstable steady-state solutions are shown in red and black, respectively. As seen in these figures, this recurrent rate-code NI (Equation 1) has only two stable attractor states, in which neurons either fire at their maximum rate (*F*_max_) or become quiescent. This implies that if there is a small perturbation in the strength of the recurrent connections or if there are changes in the sensory stimuli (e.g., a temporal gap in the incoming sensory information, *E*=0), this network could lose temporally accumulated information^11^.

The dynamics of a location-code NI relies are captured with the following equations (Equation 2):

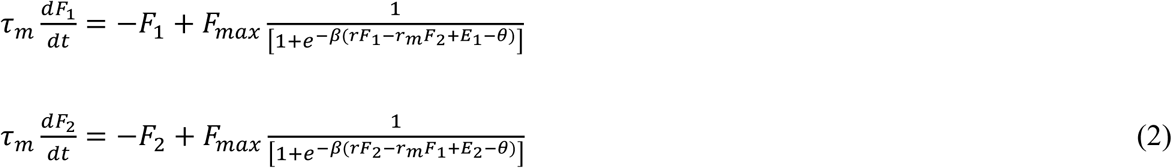

Each of the two populations had their own recurrent connections (*r*) and interacted with each other via lateral connections (*r_m_*); see Figure 3C. This mutual inhibition models the lateral inhibition mediated by SST and PV neurons in our computational model (Fig. 1). In its initial state, we assumed that population 1 fired at the maximum rate, and population 2 was quiescent; that is, population 1 had bump activity. We tested the stability of this network by examining its response to perturbations in the recurrent connections within a population (*r*), the external inputs (*E*_1_, *E*_2_) to populations 1 and 2, or the lateral interactions (*r*_m_) between the two populations.

Three main findings emerged from this analysis. First, as the recurrent-connection stregnth (*r*) increased, the network remained stable (Fig. 3D). Second, the network remained stable as we increased *E*_1_ (i.e., the external input to population 1; the red lines in Fig. 3E) but became unstable (i.e., population 1 lost its bump activity) when *E*_1_ was reduced (black lines in Fig. 3E). On the other hand, as shown in Fig. 3F, the network became unstable when *E*_2_ (i.e., the external input to population 2) increased, but it became stable when *E*_2_ decreased. In other words, the noise introduced into quiescent populations needed to be regulated in order for the network to reliably retentain information. Finally, the lateral interactions (*r_m_*) strongly impacted the stability of the network. When *r_m_* was positive but small (i.e., weak mutual inhibition), the network became unstable (the black lines in Fig. 3G). In contrast, when *r*_m_ was positive and large (i.e., strong mutual inhibition), population 1 reliably retained bump activity (Fig. 3G), and population 2 remained quiescent (Fig. 3H). That is, as long as population 1 retained bump activity intially, the mutual inhibition helped population 1 keep its bump activity. When the two populations excited each other (i.e., negative *r*_m_), neurons in both populations fired at the maximum rate (Figs. 3G and H). In this case, bump activity was not confined to population 1, indicating that a readout of bump activity based on location was not an accurate reflection of the accumulated evidence.

In brief, these analyses illustrate noticeable difference between rate- and location-code NIs. The rate-code NI encodes sensory evidence with different values of firing rate, but its steady-state response during a temporal gap is not stable (Figs. 3A and B). That is, it would lose evidence quite readily if there were even small perturbations during a temporal gap^11^. In contrast, a location-code NI is stable during a temporal gap (Figs. 3D and G), if the recurrent connecitons within a population and a mutual inhibition are sufficiently strong. Thus, during temporal gaps in sensory evidence, location-code NIs can potentially retain evidence more relibably than rate-code NIs.

### Continuous location-code neural integrator

The discrete location-code NI (Fig. 1A) has limited precision: the accumulated evidence needs to be quantized to be stored in the discrete populations. This limitation, however, is not a fundamental restriction because this discrete network can be generalized to have continuous attractor states by distributing Pyr and SST neurons into circular lattices with uniquely assigned coordinates (Fig. 1C). We call this a ‘continuous lossless integrator’. For convenience, we refer to the direction from lower to higher coordinates as the clockwise direction and higher to lower as counterclockwise. Two Pyr neurons were connected in this network if the difference between their coordinates was ≤200. Because the connections were symmetrical, each Pyr neuron made excitatory synapses with 400 of its neighboring Pyr neurons.

All Pyr and SST neurons formed non-specific connections with PV_1_ neurons. PV_2_ neurons exclusively provided feedforward inhibition to SST_1_ neurons. The connections between Pyr neurons and SST neurons were formed based on their coordinates in the circular lattice. (1) Pyr neurons made one-to-one synaptic (‘topographic’) connections with SST_1_ and SST_2_ neurons, when they had the same coordinates. (2) A SST_1_ neuron inhibited a Pyr neuron when the (absolute) difference between their coordinates was ≥200. (3) A SST_2_ neuron inhibited a Pyr neuron when the coordinate of a Pyr neuron was lower than that of a SST_2_ neuron and when the (absolute) coordinate difference was between 400 and 800. Because of this connectivity pattern, the propagation of bump activity in the counter-clockwise direction was dampened, which is possible with symmetrical chain-like recurrent connections, and only bump activity in the clockwise direction propagated through the network.

In our first analysis, we examined whether our continuous integrator could integrate sensory evidence (see Table 3 for model-parameter details). To test this integrator, we first presented a transient sensory input (time=100-200 ms) to the first 400 Pyr neurons (i.e., those with the lowest coordinates), followed by a more sustained sensory stimulus (time=100-1000) to all Pyr and PV neurons. As seen in Fig. 4A, this transient sensory stimulus elevated the rate of spiking activity strongly enough to generate bump activity. However, once generated, the feedback inhibition mediated by the PV_1_ neurons was strong enough to prevent all other excitatory neurons from spiking during the presentation of this transient sensory stimulus.

**Figure 4:**
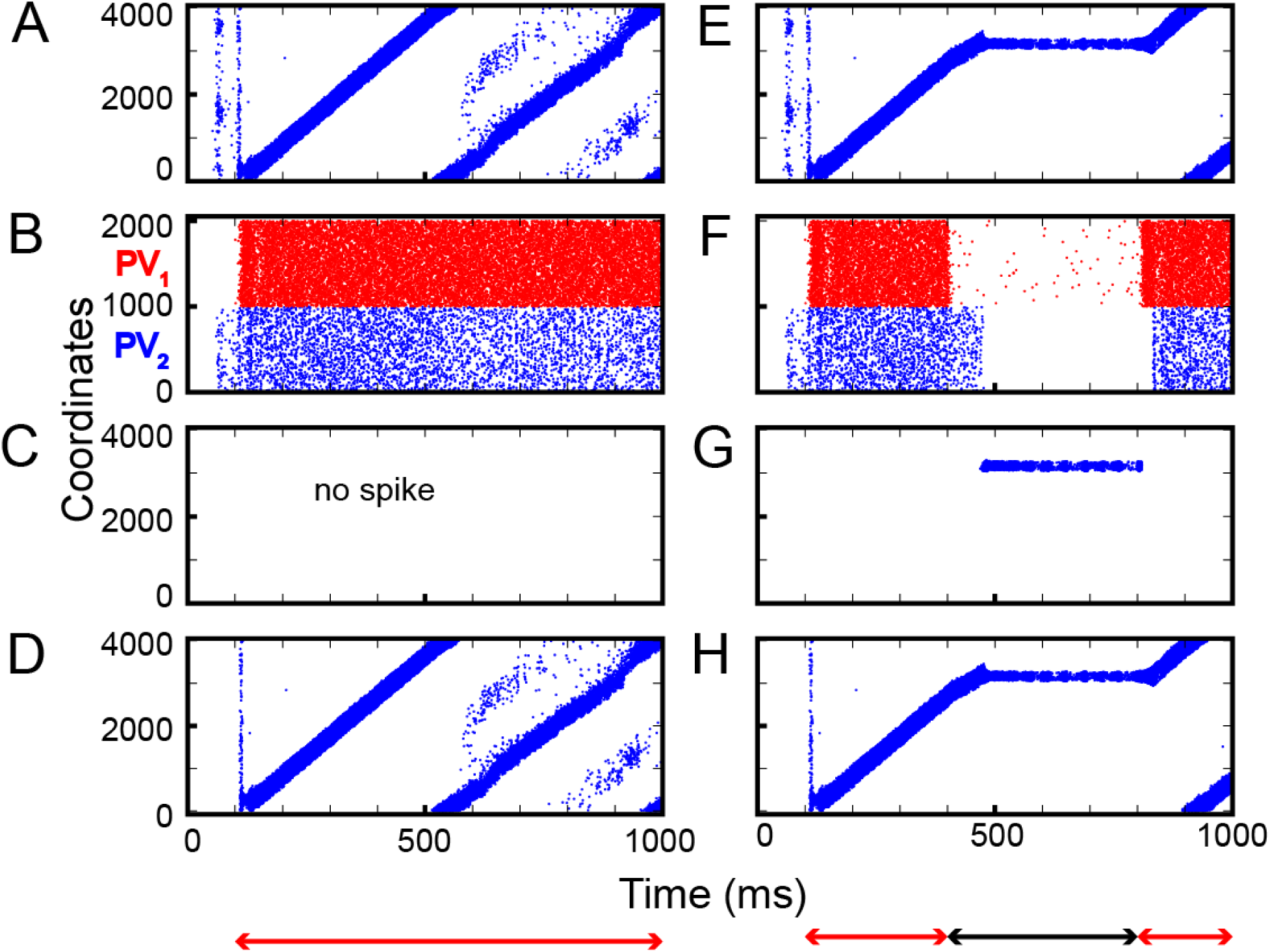
Integration of sensory inputs with and without temporal gaps. **(A)**-**(D)**, Spiking activity in Pyr, PV (PV_1_ and PV_2_), SST_1_ and SST_2_ neurons in response to constant sensory input. During stimulus presentation (100-1000 ms, marked as the red arrow), the location of bump propagates through the circular lattice: PV neurons fire asynchronously. SST_1_ neurons are quiescent, whereas SST_2_ activity mimics Pyr activity. **(E)**-**(F)**, Raster plots of Pyr, PV, SST_1_ and SST_2_ activity, respectively, when there was a temporal gap between stimulus presentations. During the gap (300-800 ms, marked by the black arrow), SST_1_ neurons became active, and the bump activity of Pyr neurons stayed at the same location.

After the offset of this transient input, bump activity propagated to other Pyr neurons in the clockwise direction (Fig. 4A). Due to the periodic boundary condition, bump activity repeatedly circulated the integrator. In our model, because excitatory synapses had not fully recovered, when the bump activity returned to the initial location, it dissipated. As a consequence, the non-specific inhibition mediated by PV_1_ neurons became weaker, which, in turn, resulted in Pyr activity at multiple locations (see Pyr cell activity after 500 ms in Fig. 4A). Concurrently, PV_1_ and PV_2_ neurons fired asynchronously (Fig. 4B). SST_1_ neurons were quiescent (Fig. 4C), but SST_2_ neurons, which received excitation from Pyr via topographic connections, mimicked Pyr activity (Fig. 4D). This SST_2_ activity prevented bump activity from propagating in the counterclockwise direction due to its asymmetrical feedback inhibition onto Pyr neurons.

Next, we tested whether this network could perform lossless integration. Like the discrete neural integrator, we presented two epochs of sensory stimuli (time=100 and 300 ms and time=800-1000 ms) that were separated by a period without sensory stimulation. For simplicity, we did not consider the onset input at 800 ms because this input had no impact on the network dynamics in the discrete integrator (Fig. 2A). As seen in Fig. 4E, bump activity cascaded through the network until there was a temporal gap in the sensory evidence. During the temporal gap, bump activity remained in the same location. Then, it resumed moving from the previous location, as information was reintroduced, consistent with lossless integration.

As in the discrete integrator, during the temporal gap in sensory information, the PV_1_ and PV_2_ neurons (Fig. 4F) became quiescent. As a result, the inhibition from the PV_1_ and PV_2_ neurons to the SST_1_ neurons was reduced, which, thereby, increased SST_1_ activity (Fig. 4G). The firing pattern of SST_2_ neurons was comparable to that of the Pyr neurons (Fig. 4H). Because the SST_1_ neurons were topographically connected to Pyr neurons, the SST_1_ inhibited non-active Pyr neurons, which prevented bump activity from propagating to a new location. Together, this transforms the network into an effective attractor network.

Next, we asked whether the dynamics of our model depended on the strength of the sensory evidence. We asked this question because neurophysiological experiments have clearly shown that the rate of accumulation of sensory evidence is positively correlated with the strength of sensory evidence, which is, subsequently, negatively correlated with reaction time^3^. In classic rate-code NIs, the firing rate increases more rapidly when sensory evidence is stronger, which readily explains the correlation between reaction-time and sensory evidence strength^1^.

In contrast, in location-code NIs, the bump location represents accumulated evidence. Thus, the propagation of bump activity would need to change as the strength of the sensory evidence changed. To address this possibility, we calculated the travel time between adjacent Pyr neurons as a function of the strength (in terms of firing rate) of the sensory inputs (evidence); the strength of the sensory evidence is controlled by α in Equation 3. Indeed, as shown in Fig. 5A, the travel time and α were inversely correlated. In other words, analogous to changes in the firing rates of rate-code NIs, as we increased the strength of the sensory evidence, bump velocity increased; examples of the propagation of bump activity through the network as a function of different values of α are shown in Fig. 5B.

**Figure 5:**
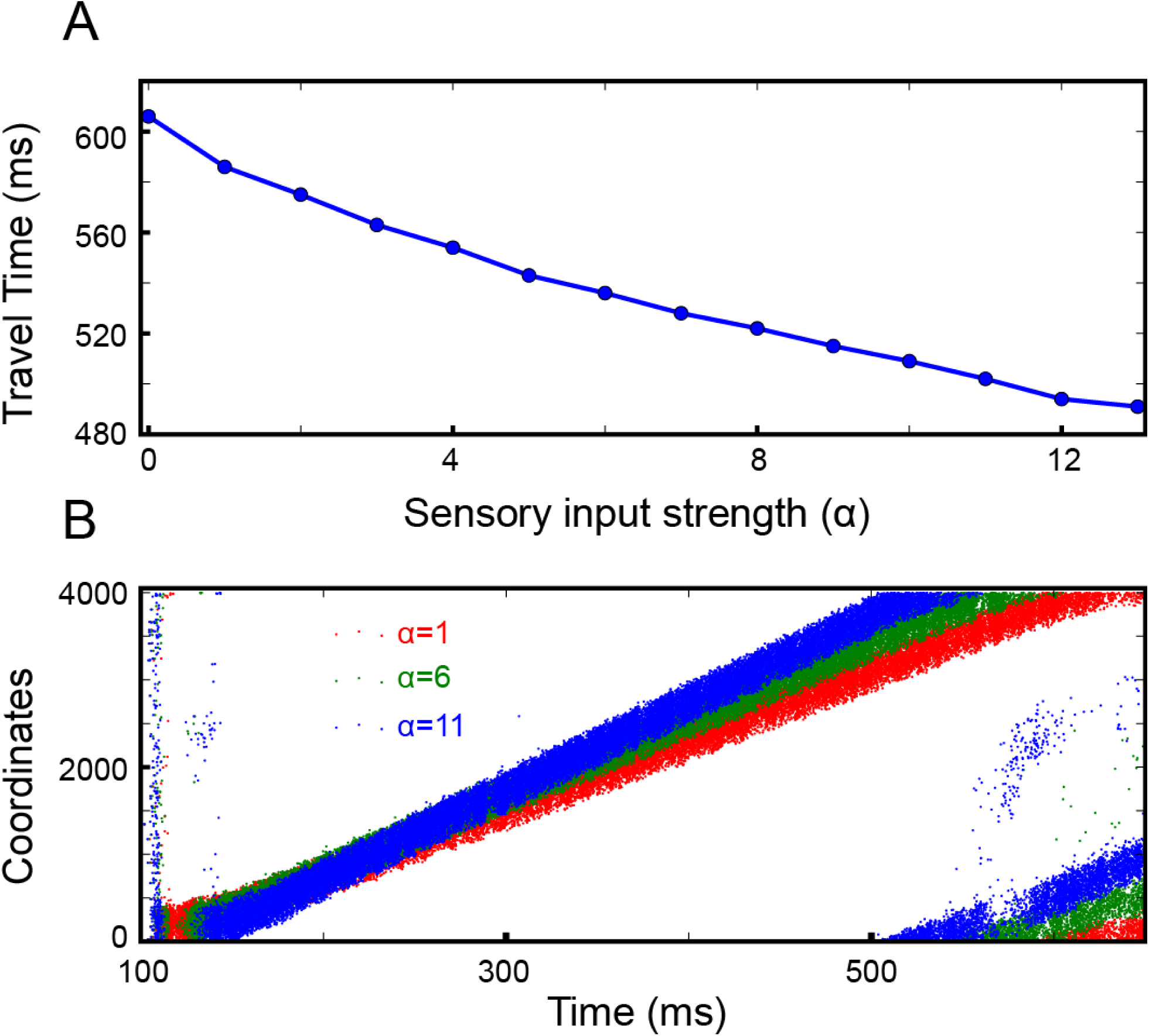
The continuous integrator was sensitive to the strength of the sensory inputs. **(A),** The travel time between consecutive Pyr neurons was inversely dependent on the strength of the sensory inputs; α represents the strength of the inputs to both Pyr and PV_1_ cells (Equation 3). **(B),** Examples of propagating bump activity as a function of different values of α.

### Decision-making with location-code NIs

Popular decision-making models, such as sequential-sampling models, suggest that a perceptual decision is made by comparing the incoming evidence to determine the most probable choice among all available choices^4, 46^. For instance, in a race model, evidence for two (or more) alternatives is accumulated until a decision threshold is reached; the alternative that reaches the boundary first would be the perceptual decision. Alternatively, evidence can be accumulated until a set time and then the alternative with the most accumulated evidence is taken as the perceptual decision. The former and latter were referred to as ‘absolute’ and ‘relative’ criterion (i.e., thresholds)^46^, respectively. Our location-code NIs can readily explain evidence integration, which suggests that these lossless integrators may be a candidate mechanism underlying race models.

Below, we propose a neural circuit that can compare the evidence accumulated in location-code NIs and produce a decision based on absolute or relative thresholds. In this work, we limit ourselves to consider two only alternatives (akin to a two-alternative forced-choice task); due to the assumption of interdependency of integrators, it is straightforward to extend the model to operate with multiple choice tasks.

#### Selective and exclusive connections between integrators and readout neurons can implement an absolute threshold for decision-making

The location of bump activity in the integrator (relative to the initial point of bump generation) can represent an absolute threshold for a single alternative. If the decision requires a comparison of two alternatives, it is necessary to find the integrator in which the bump activity arrives first at the ‘threshold’ neurons. We noted that earlier biologically-plausible models of decision-making relied on two recurrent populations with lateral inhibition^46, 47^. Although the details vary over models, in principle, two recurrent populations represent two alternatives, and the decision is represented by the exclusive activation of one of two populations; due to lateral inhibition, this exclusive activation corresponds to an attractor state in the system. Inspired by these studies, we examined if two recurrent populations, which interacted with each other via lateral inhibition, can detect the moment and the identity of integrator when the bump activity arrives first at one integrator’s threshold neurons. As seen in Fig, 6A, we built two integrators (1 and 2), each of which was connected to a distinct population of readout neurons (i.e., the recurrent network). We assumed that the last 100 (out of 4000) Pyr neurons in each integrator were ‘threshold’ neurons, and the two readout neuronal populations in the model mutually inhibited each other via di-synaptic inhibition (Fig. 6A).

**Figure 6:**
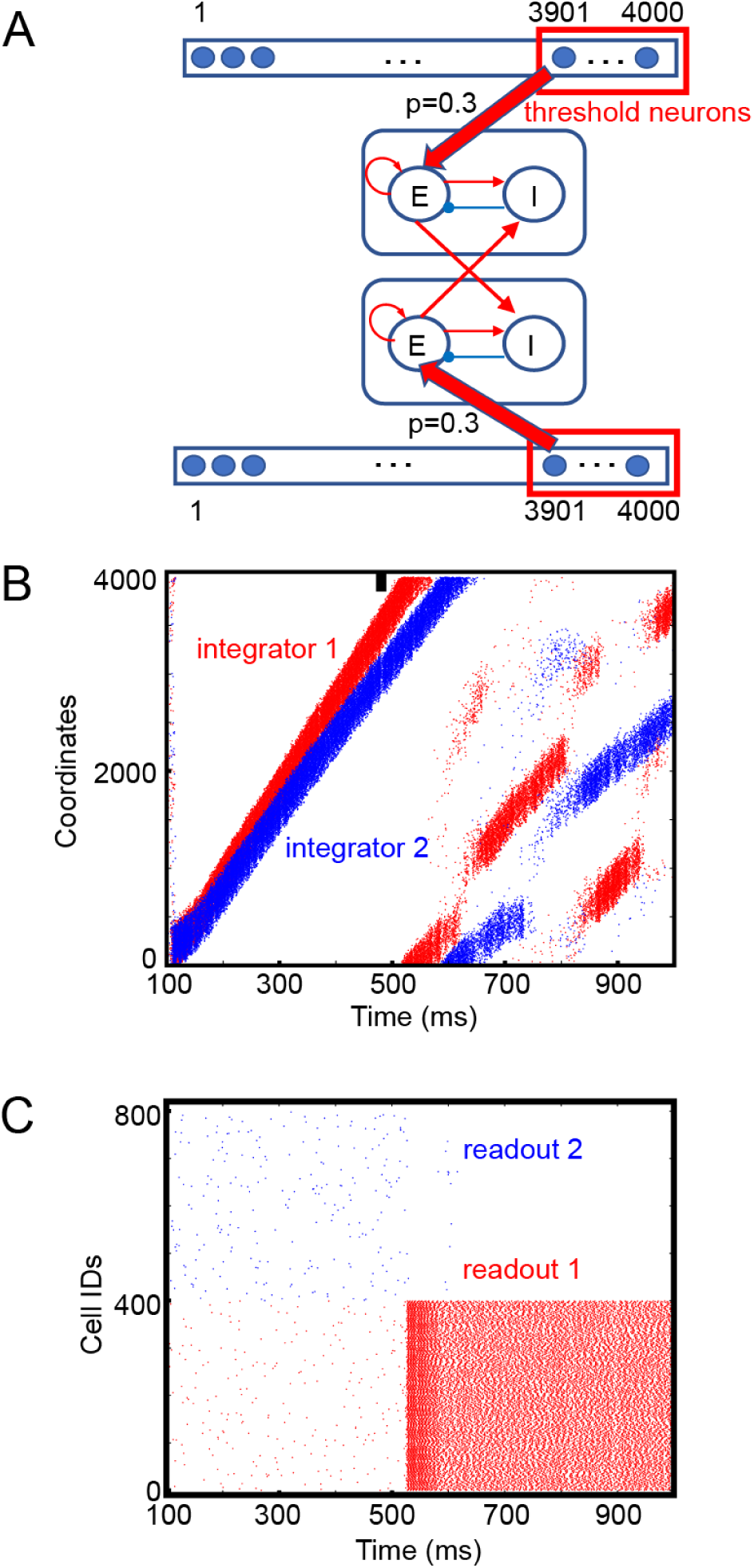
Readout schemes for decisions based on absolute thresholds. **(A)**, We assumed that there two continuous location-code integrators (top and bottom of schematic) and that 50 (out of 100) randomly chosen threshold neurons (i.e., the last 100 of the 4000 Pyr neurons in each continuous integrator) projected to excitatory neurons (E) in one of two readout neuronal populations. E neurons projected to other E and inhibitory readout neurons (I) within the population, and the connection probabilities for E-E and E-I connections were 0.3 and 0.1, respectively. The connection probabilities of cross-population connections and inhibitory connections were 0.5 and 0.1, respectively. The strengths of the excitatory and inhibitory connections were 0.12 and 0.72 pA. **(B)**, Raster plot of the two integrators. The first and second integrators are represented in red and blue, respectively. Because the first integrator had stronger stimulus inputs than the second one, bump activity propagated faster in the first integrator than in the second. The thick black vertical line represents the threshold neurons. **(C)**, Raster plots of two populations of readout neurons, shown in red and blue, respectively.

In the simulations, we titrated the strength of sensory evidence between the two integrators: integrator 1 (i.e., alternative 1) had stronger sensory inputs (α=8) than integrator 2 (i.e., alternative 2; α=3); see Equation 3. As seen Fig. 6B, the bump activity reached the threshold neurons (i.e., the last 100 Pyr neurons) faster in integrator 1 due to the stronger sensory evidence. We also noted that the readout neurons, which were exclusively connected to the threshold neurons, fired persistently even after the threshold neurons stopped firing (Fig. 6C), which was maintained via recurrent connections within the readout neurons. That is, the readout neurons not only detected the moment of crossing of threshold but also can store the decision (at least temporarily). These results suggest that the exclusive connections between the integrator and readout neurons can be a realization of an absolute threshold model of decision-making.

#### Gradient connections can implement relative thresholds for decision-making

A relative threshold requires readout neurons to track the accumulated evidence in both integrators, whenever necessary. The mechanism described above cannot track this information because the readout neurons are agnostic about the location of bump activity until it reaches the threshold neurons.

In contrast, if the readout neurons are connected to Pyr neurons in integrators via a connection probability that linearly increases as a function of the coordinates of integrator’s Pyr neurons, it is possible to realize a relative threshold. Pyr neurons in the integrator 1 projected to excitatory neurons in readout neuronal population 1 and inhibitory neurons in readout neuronal population 2. Integrator 2 is connected to readout neurons in an analogous manner (Fig. 7A). This gradient connection is consistent with the notion that synaptic connectivity (connection probability) decays over distance^48^. The maximal connection probability p_0_ in the model can determine the overall number of connections between the integrator and readout neurons.

**Figure 7:**
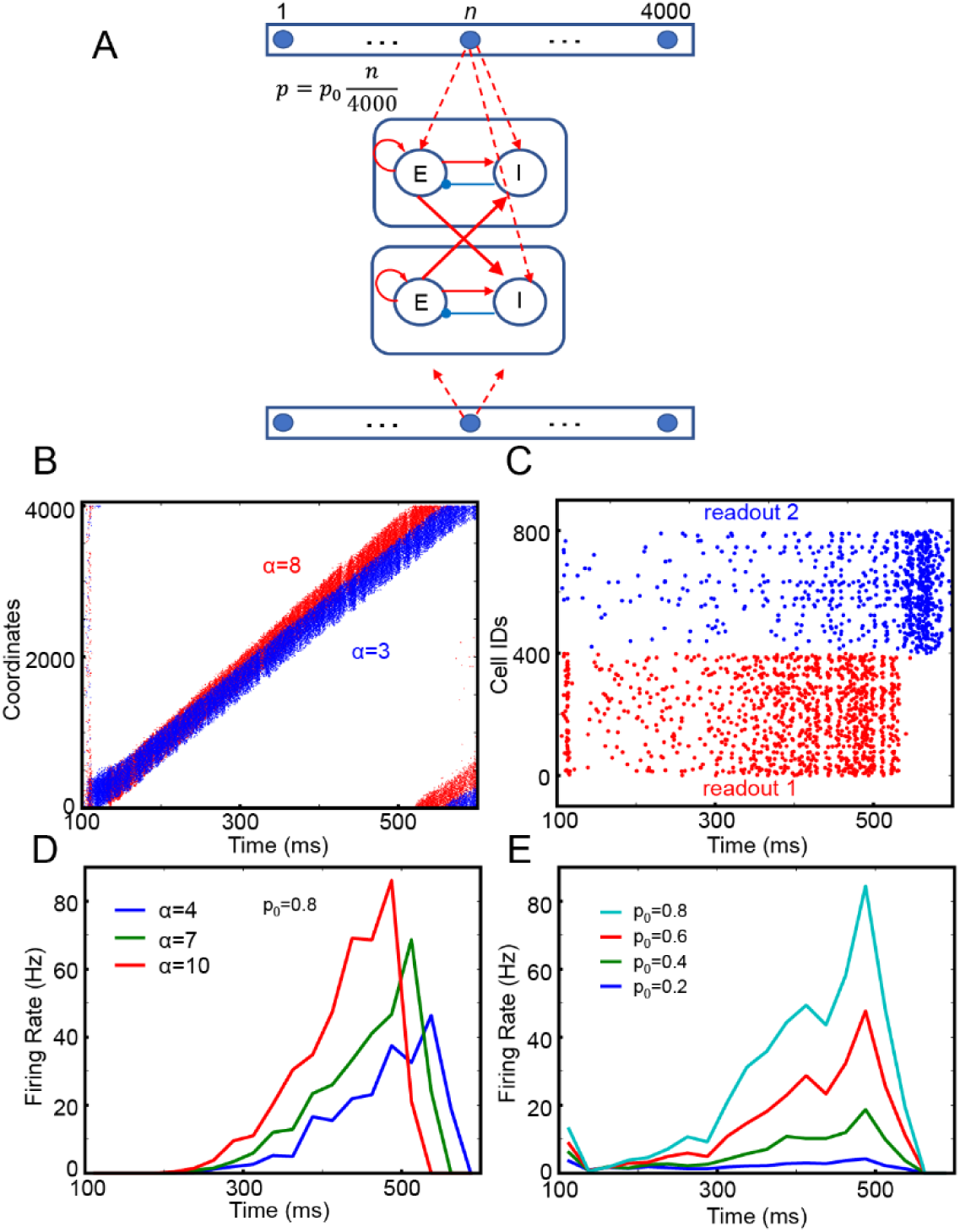
Readout schemes for decisions based on relative thresholds. **(A)**, We assumed that there two continuous integrators (top and bottom of schematic) and that each Pyr neurons in each continuous integrator projected to excitatory neurons (E) in one of two readout neuronal populations. The connection probability (p) increased, as the coordinate of Pyr neurons increased. P_0_ is the maximal connection probability. In this simulation, both E and I neurons received 200-Hz external inputs via synapses whose strength was 1.3 pA. **(B)**, Raster plot of the two integrators. The first and second integrators are represented in red and blue, respectively. Because the first integrator had stronger stimulus inputs than the second one, bump activity propagated faster in the first integrator than in the second. **(C)**, Raster plots of two populations of readout neurons, shown in red and blue, respectively. **(D)**, Time course of spiking activity of integrator-1 neurons as a function of time and the strength of sensory input to integrator 1. The strength of sensory input to integrator 2 was fixed at α=1, and p_0_ =0.8. **(E)**, Time course of spiking activity of integrator-1 neurons as a function of maximal probability p_0_.

Because integrator 1 received stronger sensory inputs (α=8) than integrator 2 (α=3), bump activity in the two integrators propagated at different speeds (Fig. 7B). As seen in Fig. 7C, readout neuronal population 1 had more activity than population 2. Further, its activity increased until bump activity returned to the initial location (due to the periodic boundary condition), suggesting that the spiking activity of readout population 1 reflected the difference between evidences. This observation is consistent with this network implementing a relative threshold for decision-making.

We also found that the spiking activity of readout population 1 is correlated with the difference in the sensory strength of sensory evidence between the two integrators (Fig. 7D). The average firing rate of readout neurons increased faster when the difference in sensory evidence between the two alternatives was stronger, which further supports our idea that this gradient-connection network can implement a relative threshold for decision-making. In addition, the spiking activity of the readout neurons in population 1 can grow more rapidly when a higher p_0_ is chosen (Fig. 7E), which is evidence that that denser connections between the integrators and the readout neurons lead to faster decision times.

#### Temporal profile of spiking activity in the readout neurons: stepping versus ramping

Rate-code NIs account for both individual and population-level ‘ramping’ (accumulating) activity in cortical regions like area LIP^2, 3, 5^. However, whereas population-level activity ramps, individual neuronal activity may be better described as ‘stepping’ activity^16, 49^.

To shed some light on the nature of these two forms of neuronal activity, we tested whether the readout neurons can reproduce either ramping or stepping activity by considering a single integrator and readout neuronal population. Specifically, we tested if single readout neuronal activity can be disassociated from population readout activity as a function of the connectivity between the integrator and the readout neuronal population.

As seen in Fig. 8A, we connected the integrator and readout neuronal population with gradient connections and varied p_0_ (the maximal probability of connections) to test the population and individual neuronal activity. When p_0_=0.1 or 1, population readout activity ramped up (Figs. 8B and C). In contrast, individual neuronal activity showed strikingly different behaviors as a function of p_0_ (Figs. 8D and E). When p_0_=0.1, individual neuronal activity did not exhibit ramping activity (Fig. 8D). However, when p_0_=1.0, individual neuronal readout activity also ramped up (Fig. 8E). To further quantify these differences in activity as a function of p_0_, we conducted a linear regression analysis between time and 25 ms-binned firing rates of individual neuronal activity. We found that when p_0_=1.0, the firing rates of most readout neurons (313 out of 400) were significantly correlated with time (p<0.05). In contrast, when p_0_=0.1, only a fraction of neurons (6 out of 400) showed significant correlation (Fig. 8F).

**Figure 8:**
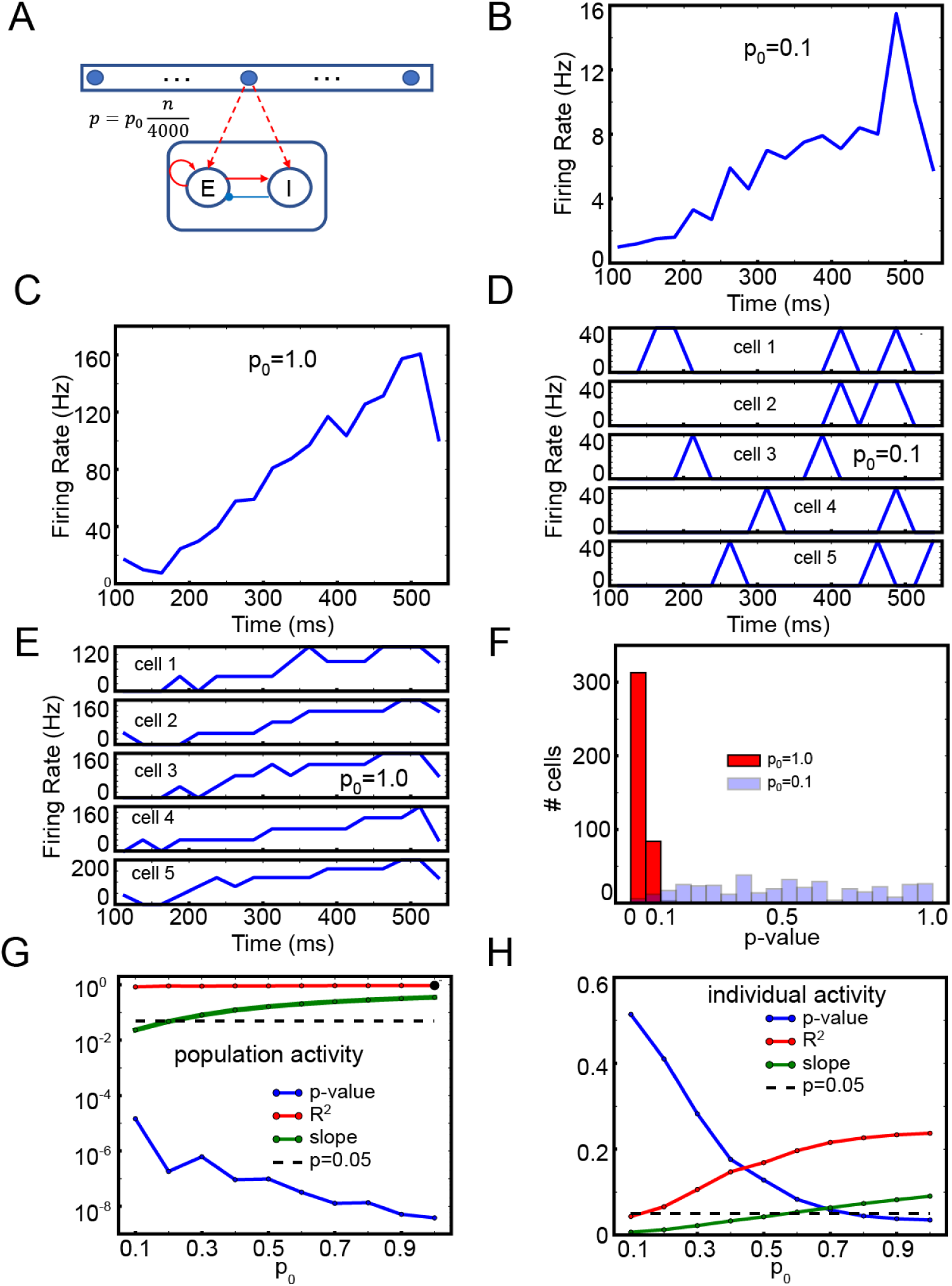
Readout neuron activity with the gradient connections. **(A)**, The structure of single set of integrator and readout neurons. **(B)**, Time course of population activity with p_0_=0.1 **(C)**, the same as (B) but with p_0_=1.0. **(D)**, Time course of individual neuro activity with p_0_=0.1 **(E)**, the same as (D) but with p_0_=1.0. **(F)**, Histograms of p-values with p_0_=0.1 and 1.0.**(G)**, Linear regression of the average firing rate of 400 *E* readout neurons depending on p_0_. **(H)**, Linear regression of individual neuron activity depending on p_0_.

We further tested a wider range of p_0_ and conducted a linear regression between population/individual neuronal activity and time. The population activity was significantly (p<0.05) correlated with the time, independent of p_0_ (Fig. 8G). As expected, the correlation between the time and individual neuron activity depended on p_0_: readout neurons produce stepping-like activity at low p_0_, but at high p_0_, we found ramping activity. When p_0_ was higher than 0.7, individual neuronal activity was significantly (p<0.05) correlated with the time (Fig. 8H).

Finally, we noticed that the individual neuronal activity was transient, unlike the experimental finding that individual cells stayed active once they stepped up to a decision^16^. However, the duration of individual neuron activity can be prolonged (supplemental Fig. 2) when the connection probability of recurrent connections in the readout neurons increased, which closer replicates this experimental finding^16^.

## Discussion

Perceptual decision-making relies on the accumulation of sensory evidence (i.e., decision-variables) that is extracted from ambiguous sensory stimuli^2, 4, 5, 46, 50–52^. It is generally thought that perceptual decision-making is instantiated through rate-code neural integrators (NIs), which are based on recurrent inputs to compensate for the leak currents^1, 8^. However, the degree to which rate-code NIs can explain perceptual decision-making is limited. For example, rate-code NIs become unstable when there is a temporal gap in the flow of incoming sensory evidence (Fig. 3), whereas behavioral studies indicate that participants act as ‘perfect/lossless’ integrators and are not affected by these temporal gaps^11, 12^.

How then can the brain make reliable decisions even with temporal gaps? We proposes that the cortex can integrate sensory evidence and maintain accumulated evidence during temporal gaps by utilizing location-code NIs, in which the location of bump activity represents the amount of presented sensory evidence^9, 10^; see below. In our simulations, bump activity in the integrator progressed through the network when sensory inputs were provided but stayed at the same location in the absence of sensory information. The location of the bump was stable due to the inhibition of SST cells (Figs. 2 and 4). This indicates that our integrator, unlike traditional rate-code NIs, can account for the robustness of perceptual decision-making during temporal gaps in sensory evidence.

We note that sequential activation, consistent with bump activity propagation in our model, has been observed in multiple brain regions^53, 54^ including the visual cortex^17–19, 55, 56^, parietal cortex^21^ and frontal cortex^57^. Notably, Harvey et al.^21^ found that posterior parietal cortex neurons were sequentially activated during decision-making, raising the possibility that location-code NIs can exist in association cortical regions like area LIP. That is, it is plausible that both location-code NIs and readout neurons coexist in area LIP, in which both stepping and ramping activity has been observed.

### Comparison to other location code NIs

In terms of function, our model reproduces the findings of previously reported location-code NIs, which modeled head-direction neurons encoding the direction of an animal’s head relative to its body and independent of its location in the environment^9^. However, the underlying mechanisms between our NI and previously described ones are quite distinct.

In previous location-code NIs, the shift in the location of bump activity was realized by so-called “rotation” neurons, which employed either strictly excitatory neurons^10^ or strictly inhibitory neurons^9^; these rotation neurons are located in the portion of the thalamus that receives inputs from the vestibular system. In contrast, we found that a cortical circuit, which consisted of excitatory pyramidal neurons and different types of inhibitory interneurons, can readily implement a location-code NI.

More specifically, two common inhibitory cortical neurons^30^ –PV and SST interneurons– made distinct contributions to this operation. PV neurons, which provided nonspecific feedback inhibition to pyramidal neurons^35, 58^, ensured that bump activity existed only at a single location. On the other hand, SST neurons mediated lateral inhibition and transformed the network into an effective attractor network capable of maintaining accumulated evidence even during temporal gaps in sensory information (Figs. 2C and 4G). We note that this theoretical finding is consistent with the empirical finding that SST cells are selectively activated during a delay period when a stimulus is removed and an animal needs to remember task-relevant information^59^. In contrast to the role that interneurons and their inhibitory synapses played in our network model, depressing excitatory synapses made bump activity propagate through the network (Figs. 2D). Together, our simulation results suggest that neurons and synapses in the neocortex are indeed suitable for controlling and maintaining the propagation of bump activity.

### Sparse and dense gradient connections may be dynamically selected depending on the demands

In our model, ramping or stepping activity can emerge depending on the afferent inputs from a location-code NI. Dense gradient connections (i.e., high p_0_) induce ramping activity. On the other hand, sparse gradient connections (i.e., low p_0_) induce stepping activity.

Dense gradient connections have a clear functional advantage: The firing rates of readout neurons increase faster and the latency of activity initiation is shorter (Fig. 7E), which could accelerate decision-making. This raises the possibility that tasks that encourage fast decisions may require dense gradient connections, which can, in turn, induce ramping activity, a classic model of decision-making.

Then, what is the functional advantage of sparse connections, which induce stepping activity in the model? Sparse connections may be optimal if decision-making is confined to a specific time window. If there is a predetermined time frame in which a decision needs to be made, it is not necessary for readout neurons to be active at all times. Instead, to reduce erroneous decisions, it may be better to suppress readout neuron activity outside the time window in which the decision needs to be made. One effective way to do this would be to lower the excitability of readout neurons and activate them only when necessary. In our model, sparse connections lowered readout neuron activity (Fig. 7E). Moreover, when external sensory inputs are introduced to readout neurons, categorical decision variables are correctly generated (supplemental Figs. 3A and B); the readout neurons can also hold the categorical decision when recurrent connections within the readout neurons are strong enough (supplemental Fig. 3C). That is, sparse connections may be used if decisions can be initiated via top-down signals such as expectation.

Together with the empirical observation^60^ that the density of synaptic connections depends on cognitive demands, we propose that stepping and ramping modes emerge from different cognitive demands or different behavioral tasks.

### Concluding remarks

Many theoretical studies have been dedicated to studying the neural correlates of persistent activity due to its potential links to cognitive functions such as decision-making^1, 8^. Recent theoretical studies have raised the possibility that the sequential activation of neurons could be the substrate of working memory^20, 56, 57^, reigniting interest in the mechanisms underlying sequential activation.

While the determination of the exact mechanisms behind any cognitive functions remains difficult, we would like to underscore that our model demonstrates that cortical circuits can natively switch between two seemingly distinct states, the stable steady state (e.g., bump activity maintenance) and the sequential activation state (e.g., bump activity propagation). We are not arguing that location-code NIs preclude the existence of rate-code Nis in neural systems. As they have distinct pros and cons, we speculate that location- and rate-code NIs are rather complementary and can be selected depending on cognitive demands.

## Methods

In this study, we developed lossless neural integrators, which were implemented within the NEST environment^63^, a peer-reviewed, freely-available simulation package. All neurons in the model were leaky integrate-and-fire (LIF) neurons. The excitatory and inhibitory neurons within an integrator formed excitatory and inhibitory connections onto a set of ‘target’ neurons. All integrator neurons and target neurons had identical internal dynamics; specifically, each presynaptic spike induced an abrupt increase in a neuron’s membrane potential that decayed exponentially. These neurons were implemented using the native NEST model iaf_psc_exp^63^. Table 1 shows the exact parameters used for the neurons and synapses in both neural integrators.

### The structure of the *discrete* integrator

The structure of the discrete integrator is summarized in Figs. 1A and B. As seen in Fig. 1A, the discrete integrator consisted of 19 different neuronal populations. 17 of these neuronal populations contained 400 pyramidal (Pyr) and 16 somatostatin (SST) model neurons. Within each of these 17 populations, Pyr neurons formed excitatory synapses with both Pyr and SST neurons. These 17 populations were topographically organized: Pyr neurons within a population had unidirectional excitatory connections with the adjacent population (e.g., population 2 projected to population 3 but not back to population 1). We had a periodic boundary condition in which the (last) population 17 connected to the (first) population 1; see Fig. 1B. In contrast, SST neurons formed inhibitory connections with Pyr neurons in all of the other populations. Recurrent connections between Pyr neurons within a particular population had depressing synapses^24–29^, but all of the other synaptic connections were static. We implemented these depressing synapses using the Tsodyks-Markram model included in the NEST distribution (Table 1).

The two remaining populations each had 1088 parvalbumin (PV) neurons. All of the Pyr neurons had excitatory connections with the PV neurons in one population (PV_1_) but not with those in the second PV population (PV_2_). Both PV_1_ and PV_2_ neurons formed non-specific inhibitory connections with Pyr and SST neurons; see Table 2 for the connection probability. These two PV populations simulated feedback and feedforward inhibition between Pyr neurons.

### The structure of the *continuous* integrator

The continuous integrator was composed of a population of Pyr neurons, two PV populations (PV_1_ and PV_2_), and two populations of SST neurons (SST_1_ and SST_2_); see Fig. 1C. Table 3 lists the parameters of these neuronal populations. In this network, 4000 Pyr, SST_1_ and SST_2_ neurons were distributed in a circular lattice, each of which had unique coordinate between 1-4000. We arbitrarily set the coordinates to increase in the clockwise direction. The neuronal numbers were arbitrary and were not constrained by the ratio of excitatory to inhibitory neurons, which is roughly 4:1. It should be noted that it is straightforward to extend this network model to include more excitatory neurons. For example, instead of a single Pyr neuron at each coordinate, a small population of Pyr neurons at each coordinate can be instantiated without changing any of the details of the network structure.

Pyr neurons were mutually connected, via excitatory connections, to their neighboring Pyr neurons when the difference between their coordinates was ≤±200, which is equivalent to a distance-dependent connection probability^48^. These connections were established with a periodic boundary condition: Pyr neuron 4000 and Pyr neuron 1 were mutually connected.

Pyr neurons interacted with the PV_1_, SST_1_ and SST_2_ populations in distinct ways. First, the pattern of connectivity between the Pyr and PV_1_ populations was randomly generated. Second, a Pyr neuron projected only to those SST_1_ and SST_2_ neurons that had the same coordinates (i.e., a one-to-one topographic mapping). The connection strength was designed to be just strong enough for a single Pyr “spike” to cause a SST_1_ or SST_2_ neuron to fire (Table 3), like a single layer-5 pyramidal-neuron spike can induce SST-expressing Martinotti neurons to fire^64^. Finally, SST_1_ and SST_2_ neurons also had inhibitory connections with Pyr neurons but had different connectivity rules. SST_1_ neurons formed connections only with those Pyr neurons in which the SST_2_-and-Pyr difference was ≥200. In contrast, SST_2_ neurons formed connections only with those Pyr neurons with lower coordinate values.

Other important model details are that PV_2_ neurons randomly inhibited SST_1_ neurons; the connection probability is shown in Table 3. Further, the PV_1_ and PV_2_ populations were independent of this circular lattice (see Fig. 1C). In our continuous integrator, all excitatory synapses were depressing, whereas all inhibitory synapses were static.

### External inputs for both integrators

The excitability of each neuron depended on the sum of its synaptic inputs from all of the other neurons in the network and from external inputs. Tables 2 and 3 show the neuron-type-specific rates of these external inputs, which were modeled with Poisson spike trains. In the model, there were ‘background’ and ‘stimulus inputs’ (i.e., sensory information). Background inputs were independent of stimulus presentations and mimicked afferent inputs from other cortex^65^. Stimulus inputs had both ‘transient’ and ‘sustained’ modes of activity. The transient mode represented the transient onsets of neural activity that have been observed in the sensory systems including retina, lateral geniculate nucleus and cortex^40–43^. We assumed that this transient activity helped to ensure that bump activity was always initiated at the same location in the network. Transient inputs (duration: 100 ms) were introduced to the first 400 and 100 Pyr neurons in the discrete and continuous integrators, respectively. In contrast, the sustained sensory inputs formed projections with all Pyr, PV_1_ and PV_2_ neurons during the entire stimulus. The frequency (I_sustained_) of the sensory inputs to PV_1_ neurons is given in Equation 3, and Pyr neurons received sensory inputs equivalent to 4×I_sustained_.

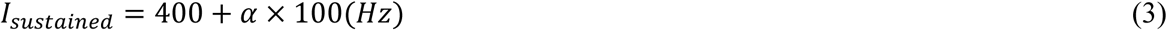

### Travelling time for the bump

Using the continuous integrator, we tested the relationship between the propagation speed of the bump and the strength of the sensory input by calculating the time course of the last 400 Pyr neurons (i.e., those with 400 highest coordinates). Specifically, we generated an event-related spike histogram using non-overlapping 10-ms bins of spiking data. ‘Travelling time of the bump’ was defined as the time, relative to stimulus onset, when the number of spikes in a single bin exceeds the sum of the mean plus two standard deviations of the number of spikes during the simulation period.

## Code availability

The simulation code is available upon request (contact JHL at) without any restrictions and will be publicly available.

## Author Contributions

JHL, JT, SV and YEC designed research; JHL performed research and analyzed data; JHL, JT, SV and YEC wrote the paper.

## Acknowledgements

JHL wishes to thank the Allen Institute founders, Paul G. Allen and Jody Allen, for their vision, encouragement and support. YEC was support by funding from the NIDCD-NIH and Boucai Hearing Restoration Fund. We also want to thank Heather Hersh and Joshua Gold for helpful comments.

## References

1. Goldman, M. S., Compte, A. & Wang, X.-J. Neural Integrator Models. Encycl. Neurosci. 6, 165–178 (2009).

2. Roitman, J. D. & Shadlen, M. N. Response of neurons in the lateral intraparietal area during a combined visual discrimination reaction time task. J. Neurosci. 22, 9475–89 (2002).

3. Gold, J. & Shadlen, M. The neural basis of decision making. Annu. Rev. Neurosci. 30, 535–574 (2007).

4. Smith, P. L. & Ratcliff, R. Psychology and neurobiology of simple decisions. Trends Neurosci. 27, 161–168 (2004).

5. Mazurek, M. E., Roitman, J. D., Ditterich, J. & Shadlen, M. N. A Role for Neural Integrators in Perceptual Decision Making. Cereb. Cortex 13, 1257–1269 (2003).

6. Kim, J. N. & Shadlen, M. N. Neural correlates of a decision in the dorsolateral prefrontal cortex of the macaque. Nat. Neurosci. 2, 176–185 (1999).

7. Ding, L. & Gold, J. I. Neural correlates of perceptual decision making before, during, and after decision commitment in monkey frontal eye field. Cereb. Cortex 22, 1052–1067 (2012).

8. Wang, X. J. Neural dynamics and circuit mechanisms of decision-making. Curr. Opin. Neurobiol. 1–8 (2012). doi:10.1016/j.conb.2012.08.006

9. Song, P. & Wang, X.-J. Angular path integration by moving ‘hill of activity’: a spiking neuron model without recurrent excitation of the head-direction system. J. Neurosci. 25, 1002–1014 (2005).

10. Skaggs, W. E., Knierim, J. J., Kudrimoti, H. S. & McNaughton, B. L. A model of the neural basis of the rat’s sense of direction. Adv. Neural Inf. Process. Syst. 7, 173–180 (1995).

11. Kiani, R., Churchland, A. K. & Shadlen, M. N. Integration of direction cues is invariant to the temporal gap between them. J. Neurosci. 33, 16483–9 (2013).

12. Liu, A. S. K., Tsunada, J., Gold, J. I. & Cohen, Y. E. Temporal Integration of Auditory Information Is Invariant to Temporal Grouping Cues 1, 2, 3. eNeuro 2, (2015).

13. Cain, N., Barreiro, A. K., Shadlen, M. N. & Shea-Brown, E. Neural integrators for decision making: a favorable tradeoff between robustness and sensitivity. J. Neurophysiol. 109, 2542–59 (2013).

14. Durstewitz, D. & Deco, G. Computational significance of transient dynamics in cortical networks. Eur. J. Neurosci. 27, 217–227 (2008).

15. Warden, M. R. & Miller, E. K. Task-Dependent Changes in Short-Term Memory in the Prefrontal Cortex. J. Neurosci. 30, 15801–15810 (2010).

16. Latimer, K. W., Yates, J. L., Meister, M. L. R., Huk, A. C. & Pillow, J. W. Single-trial spike trains in parietal cortex reveal discrete steps during decision-making. Science (80-.). 349, 182–187 (2015).

17. Beggs, J. M. & Plenz, D. Neuronal avalanches are diverse and precise activity patterns that are stable for many hours in cortical slice cultures. J. Neurosci. 24, 5216–29 (2004).

18. Ikegaya, Y. et al. Synfire Chains and Cortical Songs: Temporal Modules of Cortical Activity. Sci. (New York, NY) 559, (2004).

19. Xu, S., Jiang, W., Poo, M.-M. & Dan, Y. Activity recall in a visual cortical ensemble. Nat. Neurosci. 15, 449–55, S1-2 (2012).

20. Rajan, K., Harvey, C. D. & Tank, D. W. Recurrent Network Models of Sequence Generation and Memory. Neuron 90, 128–142 (2016).

21. Harvey, C. D., Coen, P. & Tank, D. W. Choice-specific sequences in parietal cortex during a virtual-navigation decision task. Nature 484, 62–8 (2012).

22. York, L. C. & van Rossum, M. C. W. Recurrent networks with short term synaptic depression. J. Comput. Neurosci. 27, 607–620 (2009).

23. Romani, S. & Tsodyks, M. Short-term plasticity based network model of place cells dynamics. Hippocampus 25, 94–105 (2015).

24. Markram, H., Wang, Y. & Tsodyks, M. Differential signaling via the same axon of neocortical pyramidal neurons. Proc Natl Acad Sci U S A 95, 5323–8. (1998).

25. Fuhrmann, G., Segev, I., Markram, H. & Tsodyks, M. Coding of Temporal Information by Activity-Dependent Synapses. J. Neurophysiol. 87, 140–148 (2002).

26. Reyes, A. et al. Target-cell-specific facilitation and depression in neocortical circuits. Nat. Neurosci. 1, 279–285 (1998).

27. Lefort, S. & Petersen, C. C. H. Layer-Dependent Short-Term Synaptic Plasticity between Excitatory Neurons in the C2 Barrel Column of Mouse Primary Somatosensory Cortex. Cereb. Cortex 27, 3869–3878 (2017).

28. Cheetham, C. E. J. & Fox, K. Presynaptic Development at L4 to L2/3 Excitatory Synapses Follows Different Time Courses in Visual and Somatosensory Cortex. J. Neurosci. 30, 12566–12571 (2010).

29. Petersen, C. C. H. Short-term dynamics of synaptic transmission within the excitatory neuronal network of rat layer 4 barrel cortex. J. Neurophysiol. 87, 2904–2914 (2002).

30. Rudy, B., Fishell, G., Lee, S. & Hjerling-Leffler, J. Three groups of interneurons account for nearly 100% of neocortical GABAergic neurons. Dev. Neurobiol. 71, 45–61 (2011).

31. Beierlein, M., Gibson, J. R. & Connors, B. W. Two dynamically distinct inhibitory networks in layer 4 of the neocortex. J. Neurophysiol. 90, 2987–3000 (2003).

32. Hubel, D. H. & Wiesel, T. N. Receptive fields, binocular interaction and functional architecture in the cat’s visual cortex. J. Physiol. 160, 106–154.2 (1962).

33. Hubel, D. H. & Wiesel, T. N. Receptive fields and functional architecture of monkey striate cortex. J.Physiol.(London) 195, 215–243 (1968).

34. Ko, H. et al. The emergence of functional microcircuits in visual cortex. Nature 496, 96–100 (2013).

35. Ma, W. -p. et al. Visual Representations by Cortical Somatostatin Inhibitory Neurons--Selective But with Weak and Delayed Responses. J. Neurosci. 30, 14371–14379 (2010).

36. Markram, H. et al. Interneurons of the neocortical inhibitory system. Nat. Rev. Neurosci. 5, 793–807 (2004).

37. Zhang, S. et al. Long-range and local circuits for top-down modulation of visual cortex processing. Science (80-.). 345, 660–665 (2014).

38. Adesnik, H., Bruns, W., Taniguchi, H., Huang, Z. J. & Scanziani, M. A neural circuit for spatial summation in visual cortex. Nature 490, 226–31 (2012).

39. . Jiang, X. et al. Principles of connectivity among morphologically defined cell types in adult neocortex. Science (80-.). 350, aac9462-aac9462 (2015).

40. Cleland, B. G., Dubin, M. W. & Levick, W. R. Sustained and transient neurones in the cat’s retina and lateral geniculate nucleus. J. Physiol. 217, 473–496 (1971).

41. Piscopo, D. M., El-Danaf, R. N., Huberman, A. D. & Niell, C. M. Diverse Visual Features Encoded in Mouse Lateral Geniculate Nucleus. J. Neurosci. 33, 4642–4656 (2013).

42. De Valois, R. L., Cottaris, N. P., Mahon, L. E., Elfar, S. D. & Wilson, J. A. Spatial and temporal receptive fields of geniculate and cortical cells and directional selectivity. Vision Res. 40, 3685–3702 (2000).

43. de la Rocha, J., Marchetti, C., Schiff, M. & Reyes, A. D. Linking the Response Properties of Cells in Auditory Cortex with Network Architecture: Cotuning versus Lateral Inhibition. J. Neurosci. 28, 9151–9163 (2008).

44. Ermentrout, G. B. & David, H. T. Mathematical Foundation of Neuroscience. (springer, 2010).

45. Ermentrout, B. XPPAUT. Scholarpedia 2, 1399 (2007).

46. Ratcliff, R. & Smith, P. L. A Comparison of Sequential Sampling Models for Two-Choice Reaction Time. Psychol Rev. 111, 333–367 (2004).

47. Wang, X. J. Neural dynamics and circuit mechanisms of decision-making. Curr. Opin. Neurobiol. 1–8 (2012). doi:10.1016/j.conb.2012.08.006

48. Perin, R., Berger, T. K. & Markram, H. A synaptic organizing principle for cortical neuronal groups. Proc. Natl. Acad. Sci. U. S. A. 108, 5419–24 (2011).

49. Miller, P. & Katz, D. B. Stochastic transitions between neural states in taste processing and decision-making. J. Neurosci. 30, 2559–2570 (2010).

50. Miller, P. Decision Making Models. Encyclopedia of Computational Neuroscience (Springer-Verlag New York, 2015). doi:10.4249/scholarpedia.1448

51. Ratcliff, R. A theory of memory retrieval. Psychol. Rev. 85, 59–108 (1978).

52. LaBerge, D. A recruitment theory of simple behavio. Psychometrika 27, 375–396 (1962).

53. Tang, A. et al. A Maximum Entropy Model Applied to Spatial and Temporal Correlations from Cortical Networks In Vitro. J. Neurosci. 28, 505–518 (2008).

54. Pulvermuller, F. & Shtyrov, Y. Spatiotemporal signatures of large-scale synfire chains for speech processing as revealed by MEG. Cereb. Cortex 19, 79–88 (2009).

55. Sato, T. K., Nauhaus, I. & Carandini, M. Traveling Waves in Visual Cortex. Neuron 75, 218–29 (2012).

56. Beggs, J. M. & Plenz, D. Neuronal avalanches in neocortical circuits. J. Neurosci. 23, 11167–77 (2003).

57. Seidemann, E., Meilijson, I., Abeles, M., Bergman, H. & Vaadia, E. Simultaneously recorded single units in the frontal cortex go through sequences of discrete and stable states in monkeys performing a delayed localization task. J. Neurosci. 16, 752–768 (1996).

58. Bock, D. D. et al. Network anatomy and in vivo physiology of visual cortical neurons. Nature 471, 177–182 (2011).

59. Kim, D. et al. Distinct Roles of Parvalbumin- and Somatostatin-Expressing Interneurons in Working Memory. Neuron 92, 902–915 (2016).

60. Purushothaman, G. & Bradley, D. C. Neural population code for fine perceptual decisions in area MT. Nat. Neurosci. 8, 99–106 (2005).

61. Goldman, M. S. Memory without Feedback in a Neural Network. Neuron 61, 621–634 (2009).

62. Lundqvist, M. et al. Gamma and Beta Bursts Underlie Working Memory. Neuron 90, 152–164 (2016).

63. Gewaltig, M.-O. & Diesmann, M. NEST (NEural Simulation Tool). Scholarpedia 2, 1430 (2007).

64. Silberberg, G. & Markram, H. Disynaptic inhibition between neocortical pyramidal cells mediated by Martinotti cells. Neuron 53, 735–46 (2007).

65. Potjans, T. C. & Diesmann, M. The cell-type specific cortical microcircuit: relating structure and activity in a full-scale spiking network model. Cereb. Cortex 24, 785–806 (2014).

