## Supplementary Materials for "Cortical circuit-based lossless neural integrator for perceptual decision-making"

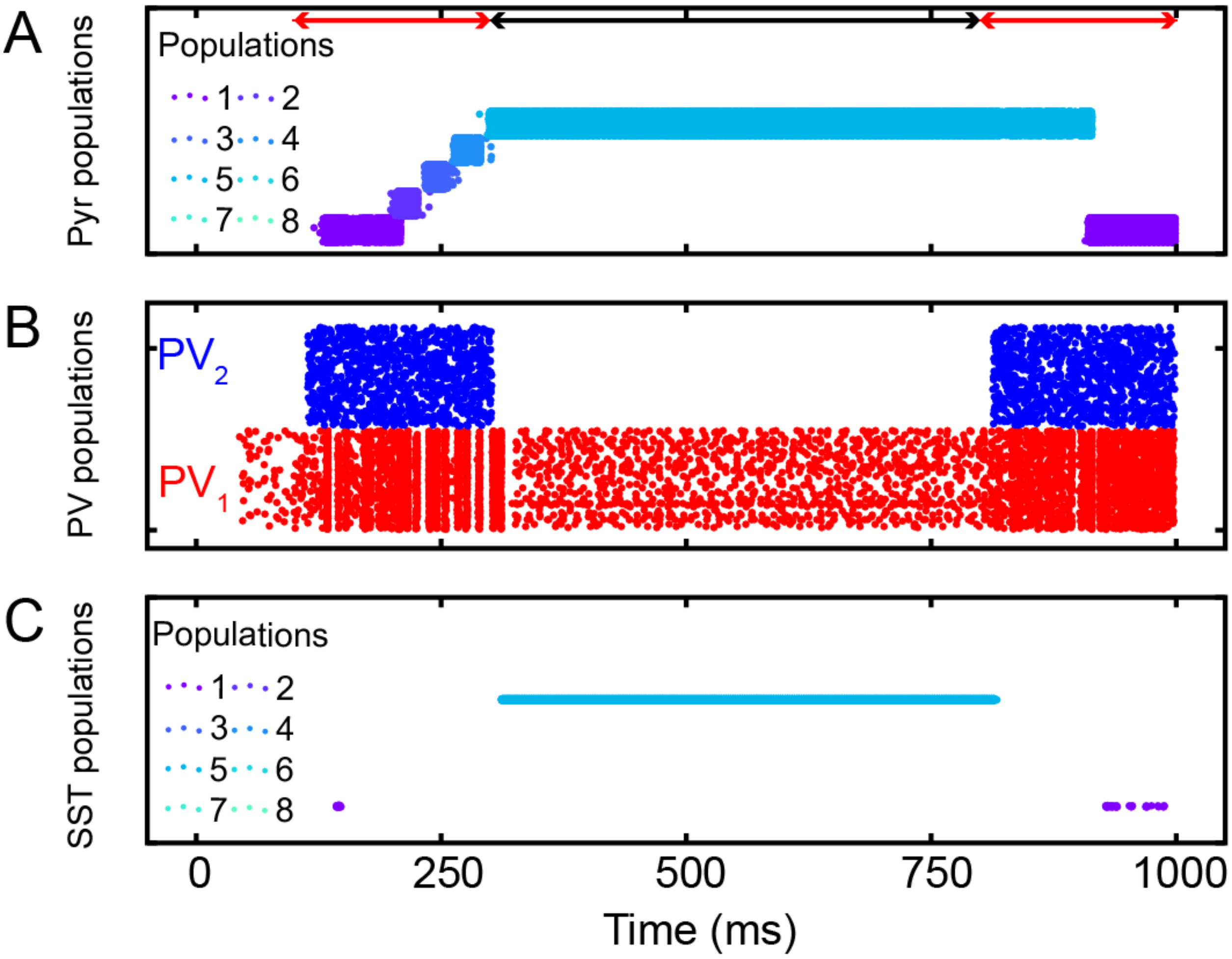


**Supplemental Figure 1: The responses of populations without inhibition of $PV_1$ to $Pyr_1$ populations during the temporal gap.** We repeated the same experiment shown in Fig. 3, but removed inhibition from PV1 to Pyr1 populations at 500 ms. That is, the first half of the responses are identical to those shown in Fig. 3. But the last half are different. As expected, when stimulus inputs were reintroduced, population 1 was reactivated by the transient inputs; this destroyed the accumulated information. Conversely, the feedback inhibition of $PV_1$ neurons ensures the network accurately encodes the evidence accumulated.

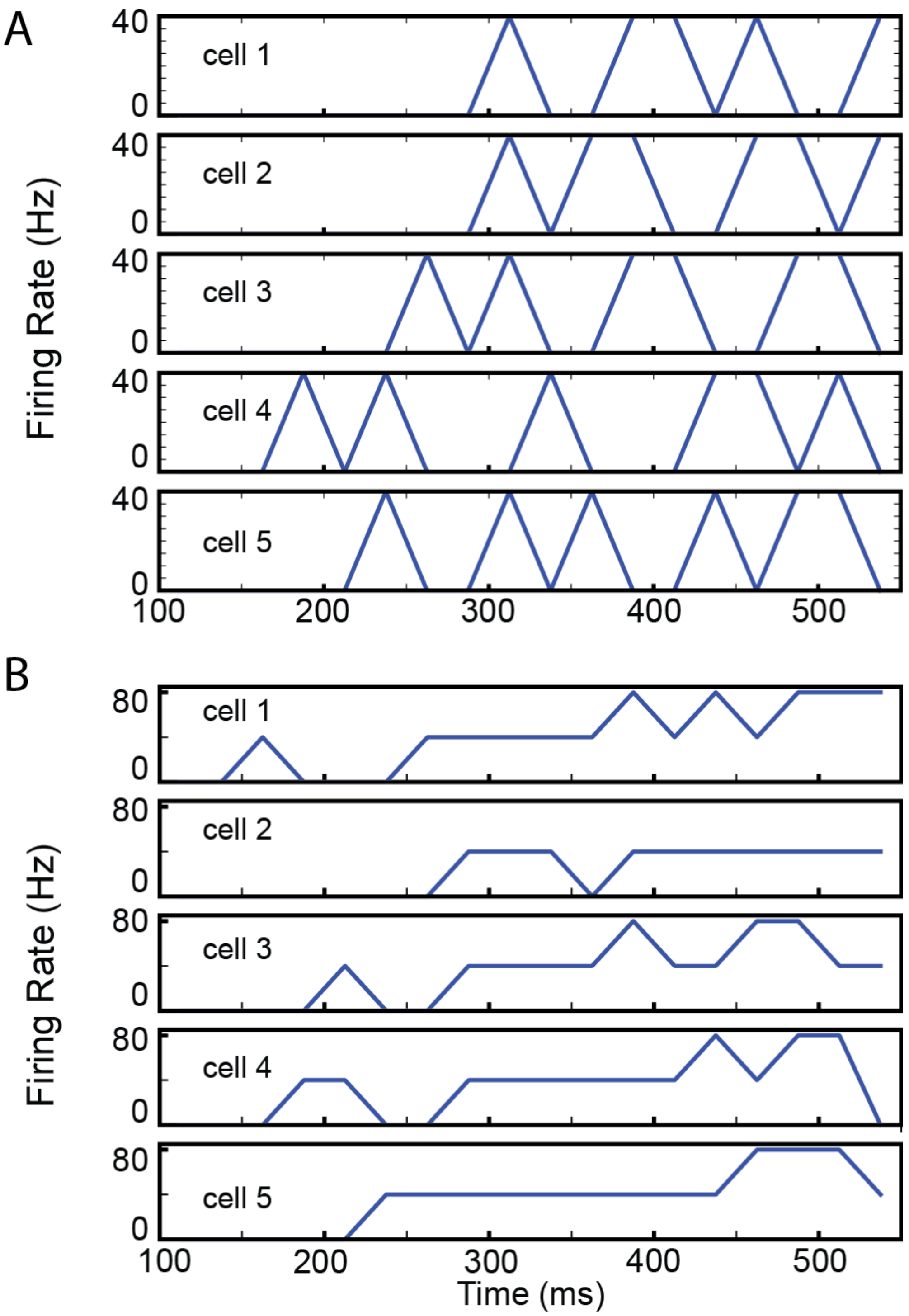


**Supplemental Figure 2: Individual readout neurons with gradient connections.** This figure illustrates the individual readout neurons with gradient connections, as in Fig. 8D. All parameters chosen are the same as those in Fig. 8D but with more number of recurrent connections among *E* readout neurons. The default connection probability ($P_{conn}$) is 0.1 (Fig. 8G). **(A)**, Individual readout neuron activity with $P_{conn}$=0.2. **(B)**, Individual readout neuron activity with $P_{conn}$=0.25.

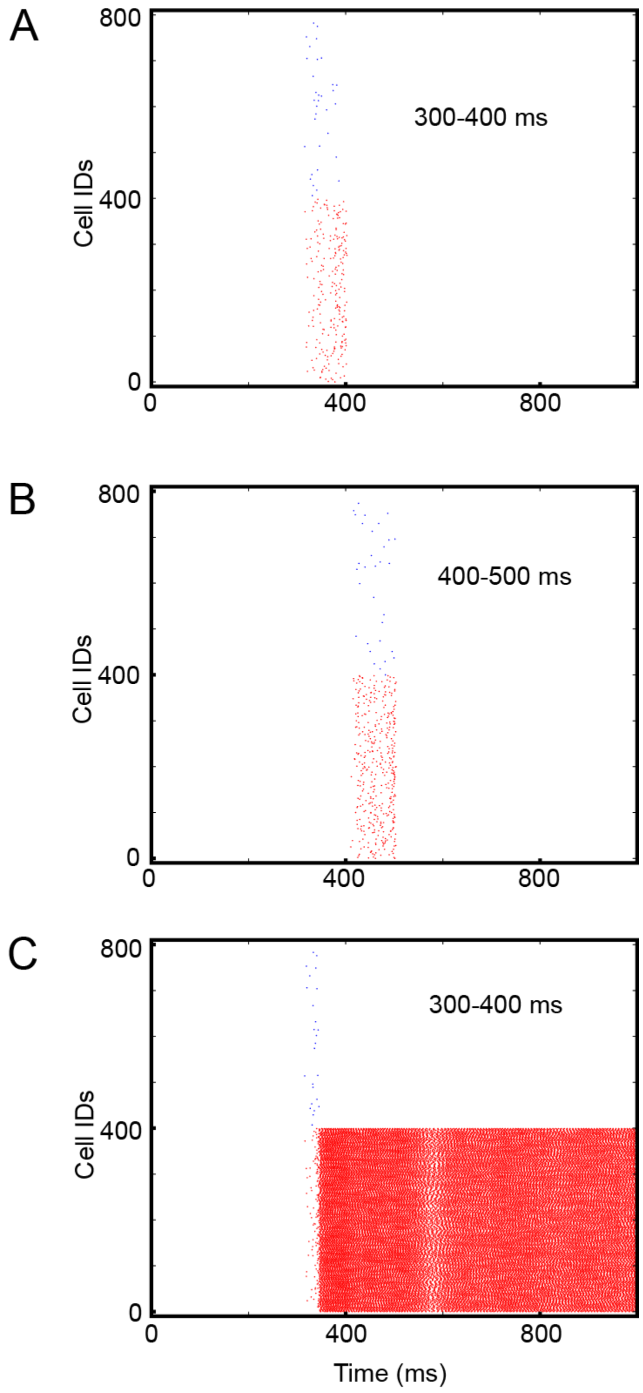


**Supplemental Figure 3: Readout neuron activity with sparse gradient connections. (A)**, Readout neuron responses. Spikes from readout neuron population 1 are shown in red, whereas those from readout neuron population 2 are shown in blue. The external inputs to excitatory neurons are introduced between 300-400 ms. The structure of readout neurons is the same as those shown in Fig. 7A, and inhibitory readout neurons receive external inputs throughout the simulation. **(B)**, the same as with (A) with external inputs to excitatory neurons between 400-500 ms. **(C)**, the same as (A) but with higher connection probability among excitatory readout neurons are increased from 0.1 to 0.38.
